# *Drosophila* neural stem cells are polarized by their daughter cells

**DOI:** 10.1101/182592

**Authors:** Nicolas Loyer, Jens Januschke

**Affiliations:** Cell & Developmental Biology, School of Life Sciences, University of Dundee, Dundee, UK

## Abstract

Controlling the orientation of cell division is important in the context of cell fate choices and tissue morphogenesis. However, the mechanisms providing the required positional information remain incompletely understood. Here we use stem cells of the *Drosophila* larval brain that stably maintain their axis of polarity and division between cell cycles to identify cues that orient cell division. Using live cell imaging of cultured brains, laser ablation and genetics we reveal that these cells use the position of their last-born daughter cell as a polarizing cue. Remarkably, this daughter cell derived signal received at one pole of the stem cell has an effect on the opposite pole influencing where apical will be in the next mitosis, thereby directing the orientation of division. Therefore, in addition to known intrinsic cues, stem cells in the developing fly brain are polarized by an extrinsic signal that acts upstream of apico-basal polarity establishment.

## Introduction

The orientation of cell division is important for cell fate choices and impacts on the morphology and function of tissues ^1-3^. Therefore, perhaps not surprising, defects in spindle orientation have been linked to developmental defects and diseases ^4,5^. Coupling spindle orientation to unequal segregation of fate determinants is one strategy during asymmetric cell division to generate different cell fates ^6,7^. Spindle orientation also affects the placement of daughter cells after division. This can alter function and fate of the resulting daughter cells as the microenvironment in different positions can cause daughter cells to experience different signals ^8^.

In cells, an evolutionary conserved molecular machinery helps to position the spindle by anchoring the astral microtubules to cortical attachment sites ^9-11^. A key challenge in this context is understanding the spatial information that determines the position of these attachment sites. The machinery anchoring microtubules at the cortex frequently depends on the axis of polarity of the dividing cell. In those contexts, the symmetry breaking event that polarizes a cell and gives the polarity axis its orientation, also determines the orientation of the subsequent division. Microtubules can act in many contexts as such a signal biasing with which orientation cells polarize (reviewed in ^12^), but a variety of other polarizing cues exist that polarize cells and orient their division.

Embryonic neural stem cells (neuroblasts) in *Drosophila* for instance can use spindle microtubules to deliver components of the microtubule anchoring machinery to the cortex ^13^. These cells can also read extrinsic cues, orienting their division perpendicular to the overlying epithelium ^14^. This is mediated by G-protein coupled receptor signalling recruiting factors directly orienting the spindle towards this signal ^15^. In other contexts, E-cadherin (E-Cad) rich cell-cell adhesion sites provide spatial information to orient the mitotic spindle ^16-19^. In the case of *C. elegans,* the site of sperm entry defines anterior posterior polarity and the orientation of the first division ^20^. The midbody resulting from this division is further used as a spatial cue orienting the subsequent P1 cell division ^21^. Cytokinesis is also linked to division orientation control in budding yeast, where the orientation of future divisions is biased by a landmark at the site of abscission ^22^. Therefore, positional landmarks linked to cytokinesis can control the orientation of cell division.

An ideal system to study mechanisms that orient cell division are the highly proliferative neuroblasts of the *Drosophila* larva that divide over many cell cycles with very little deviation in the orientation of division between different cycles. The mechanisms controlling this process are only partially understood. In neuroblasts, cortical polarity is established by the activity of the Par complex ^23-27^. The Pins (*Drosophila* homolog of LGN) complex ^28-32^ then couples the orientation of the mitotic spindle with apico-basal polarity, such that both are aligned. Interestingly, after each division the polarized localization of both complexes on the neuroblast cortex is lost, but reforms with the same orientation in the next mitosis ^33-35^. Contrary to embryonic neuroblasts ^14^, this occurs regardless of whether larval neuroblasts reside within the brain or are in isolation in primary culture ^36-38^. Currently, this process is believed to occur through the apically localized centrosome and microtubules, which act as cell intrinsic polarizing cues ^39^. However, disruption of theses cues, either through depolymerisation of microtubules or mutation in *sas4* leading to loss of centrioles ^40^, only results in a partial defect of division orientation maintenance ^37^. This suggests other polarizing cues contribute in parallel to maintain the orientation of the axis of neuroblast division.

Given that cytokinesis-related cues can direct spindle orientation in other cell types, we hypothesised that neuroblasts could use a spatial cue provided by their last-born daughter cell to orient cell division in the subsequent mitosis. Indeed, we found that neuroblasts align their divisions with the position of the last-born daughter cell (called ganglion mother cell, GMC). Disruption of the integrity of the neuroblast/GMC interface, either through laser ablation or by depletion of proteins specifically localizing to this interface, including the midbody and midbody-associated structures, perturbs neuroblast division orientation memory while it does not affect alignment of the mitotic spindle with cortical polarity. Thus, the last-born GMC is an extrinsic polarizing cue for larval brain neuroblasts in *Drosophila* orienting their axis of polarity and consequently division.

## Results

### The division axis of neuroblasts follow GMCs movements

To test our hypothesis that the orientation of neuroblast division is under the influence of extrinsic cues provided by their daughter cells, we analysed the relationship of the orientation of neuroblast division with the position of the last-born daughter cell (GMC). We used confocal imaging of whole mount brains to capture the entire volume of neuroblasts over several rounds of divisions (**VIDEO 1**), and developed a method to measure the deviation of their division axis in 3D using the characteristic shape of telophase neuroblasts as reference (**Figure 1A** and **S1A**). Using this method, we reproduced the previous observation ^37^ that maintenance of the division axis of neuroblasts is affected, but not abolished, in *sas4* mutants (Figure S1B-B’’). This partial maintenance of the division axis upon loss of the *sas4* dependent polarity cue is consistent with the continued activity of an additional polarizing cue.

As previously reported ^37^, the division axis of control neuroblasts is not perfectly maintained between cell cycles (**Figure B-C**). We next reasoned that, if the GMC provides a polarizing cue, the division axis of neuroblasts should align with the position of the GMC when neuroblasts polarize, i.e. when they start rounding at the onset of mitosis (Figure S1C, **VIDEO 2**). Thus, we tracked the position of the last-born GMC until neuroblasts started rounding up, at which point we defined a neuroblast-GMC axis (Figure 1B, magenta arrow). Comparing this axis to the following division axis (**Figure 1B**, green arrow) revealed that the position of the last-born GMC at the onset of NB rounding predicted significantly better the orientation of the subsequent division than the previous axis of division (**Figure 1C**).

We further observed that in cases for which the division axis was most highly misaligned with the previous one, the last-born GMC moved away from its initial birthplace and the neuroblast from which it derived, re-aligned its subsequent division with the new position of that GMC (**Figure 1A**). Importantly, this displacement of the GMC and the subsequent realignment of the division axis was not caused by a rotation of the entire brain, nor of the [**fig1]** neuroblast-progeny cluster as only the last-born GMC and not its neighbouring cells significantly changed position (**Figure S1D-D’’**). Whether such GMC movements are physiologically relevant or experimentally induced is unclear. Nonetheless, our observation that neuroblast divisions realign with displaced GMCs supports the idea that the GMC is a spatial cue that orients neuroblasts divisions.

### Ablation of the last-born GMC disrupts the maintenance of the orientation of neuroblast division

If GMCs provide a polarizing cue, their ablation should affect the maintenance of the axis of neuroblast division. We tested this idea by observing a neuroblast division and destroying the GMC born from this division by biphotonic laser-mediated ablation. We then observed the following division and measured its alignment with the previous one. Neuroblasts immediately deformed toward the destroyed GMC following its ablation (**Figure S2, VIDEO 3**). Therefore, we were concerned that this deformation and cellular debris generated close to the neuroblast together with damage caused by the laser could affect the maintenance of the division axis and bias our analysis. To account for such effects, we performed “control ablations” by ablating other cells in contact with the neuroblast, away from the last-born GMC (**Figure 2 A, A’**).

Control ablations also resulted in neuroblast deformation towards the destroyed cell (**Figure S2, VIDEO 4)**. However, they did not affect the maintenance of neuroblast division orientation. In contrast, ablation of the last-born GMC significantly affected this process (**Figure 2A’’, VIDEO 5**). Importantly, neuroblasts that misoriented their division following ablation and then divided again, aligned this third division with the previous, misoriented division (**Figure 2BB’’**). Thus, ablation of the last-born GMC results in a transient defect in the orientation of the neuroblast division axis, that is restored upon the generation of a new GMC. This is further consistent with our (Figure 1), and previous observations that neuroblasts reorient their division axis to align with displaced GMCs ^37^.

**Figure 1:**
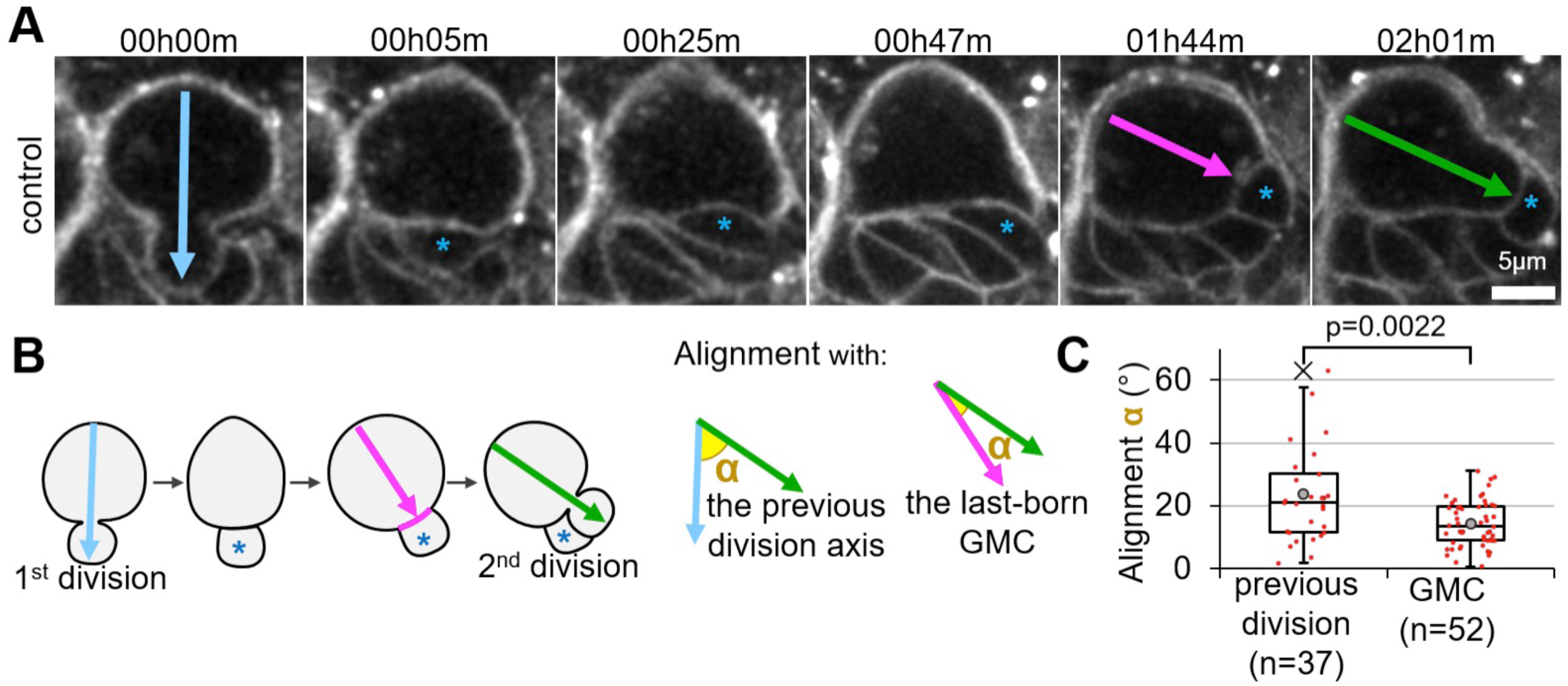
Neuroblasts realign their division axis with relocalized GMCs. **// A)** Two successive divisions of a larval neuroblast expressing the membrane marker PH::RFP. Arrows: division axis. Asterisk: last-born GMC. **B)** Angles α measured in the previous panel and quantified in the next panel. The alignment of the second division (green arrow) with the previous one (blue arrow) and with an axis (magenta arrow) defined by the GMC (blue asterisk)/neuroblast interface (magenta line) at prophase was measured in 3D. **C)** Distribution of the angles α described in the previous panel: alignment of the second division axis with theprevious one and the GMC (14±7°, n=37 vs. 24±15°, n=52). In this boxplot and every following one: cross: maximum outlier; grey circle: average; red dots: individual measurements. **//**

**Figure 2:**
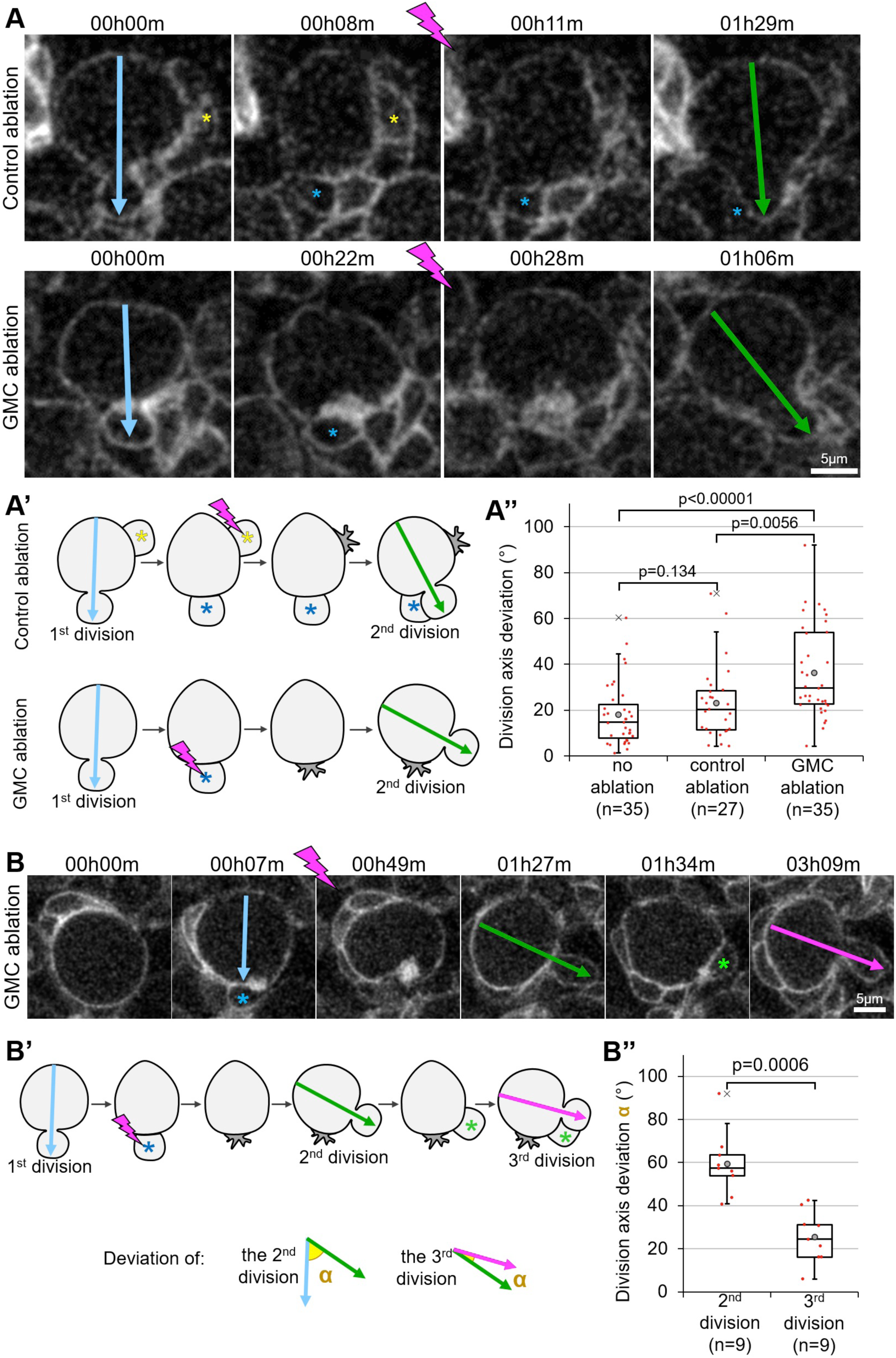
GMC ablation disrupts division axis maintenance. **// A)** Two successive divisions of neuroblasts expressing *worniu*-GAL4-driven PH::GFP pre and post ablation are shown. Upper panels: control ablation of a neuroblast-neighboring cell (yellow asterisk) away from the last-born GMC (blue asterisk). Lower panels: GMC ablation. The ablation takes place between the two frames separated by a lightning symbol. Arrows: division axis. **A’)** Schematic of the ablations and angle measurements. **A’’)** Deviation of the division axis following no ablation (Average angle: 18±14°, n=35), control ablation (23±16°, n=27) or GMC ablation (36±20°, n=35; *also shown in* ***Figures 3B*** *and* ***5C***). **B)** Three successive divisions of neuroblasts expressing *worniu*-GAL4-driven PH::GFP. The ablation takes place between the two frames separated by a lightning symbol. Blue asterisk: GMC prior to ablation, green asterisk: new born GMC. Arrows: division axis. **B’)** Schematic of the angles α measured from the movies shown in the previous panel. **B’’)** Deviation α of the second (59±22°, n=9) and third (25±11°, n=9) divisions described in the two previous panels. **//**

### The last-born GMC provides an extrinsic cue for neuroblast polarization

In neuroblast the spindle is aligned with the apico-basal polarity axis. Altered division orientation caused by GMC ablation could therefore be the result of misalignment of the mitotic spindle with the apico-basal polarity axis. Alternatively, the orientation of the polarity axis itself could be affected. We tested this by repeating GMC ablation experiments in neuroblasts expressing the apical polarity marker Baz::GFP, together with the centriole marker Asl::YFP to visualize the mitotic spindle poles. Following GMC ablation, every case of division axis misalignment (n=7 cases with deviation > 45°) displayed a misplaced apical crescent with which the spindle properly aligned (**Figure 3A-A’**). Thus, defective division axis maintenance upon GMC ablation results from altered orientation of the polarity axis, and not from defects in downstream mechanisms related to spindle anchoring. We conclude that the last-born GMC is a polarizing cue for larval brain neuroblasts.

**Figure 3:**
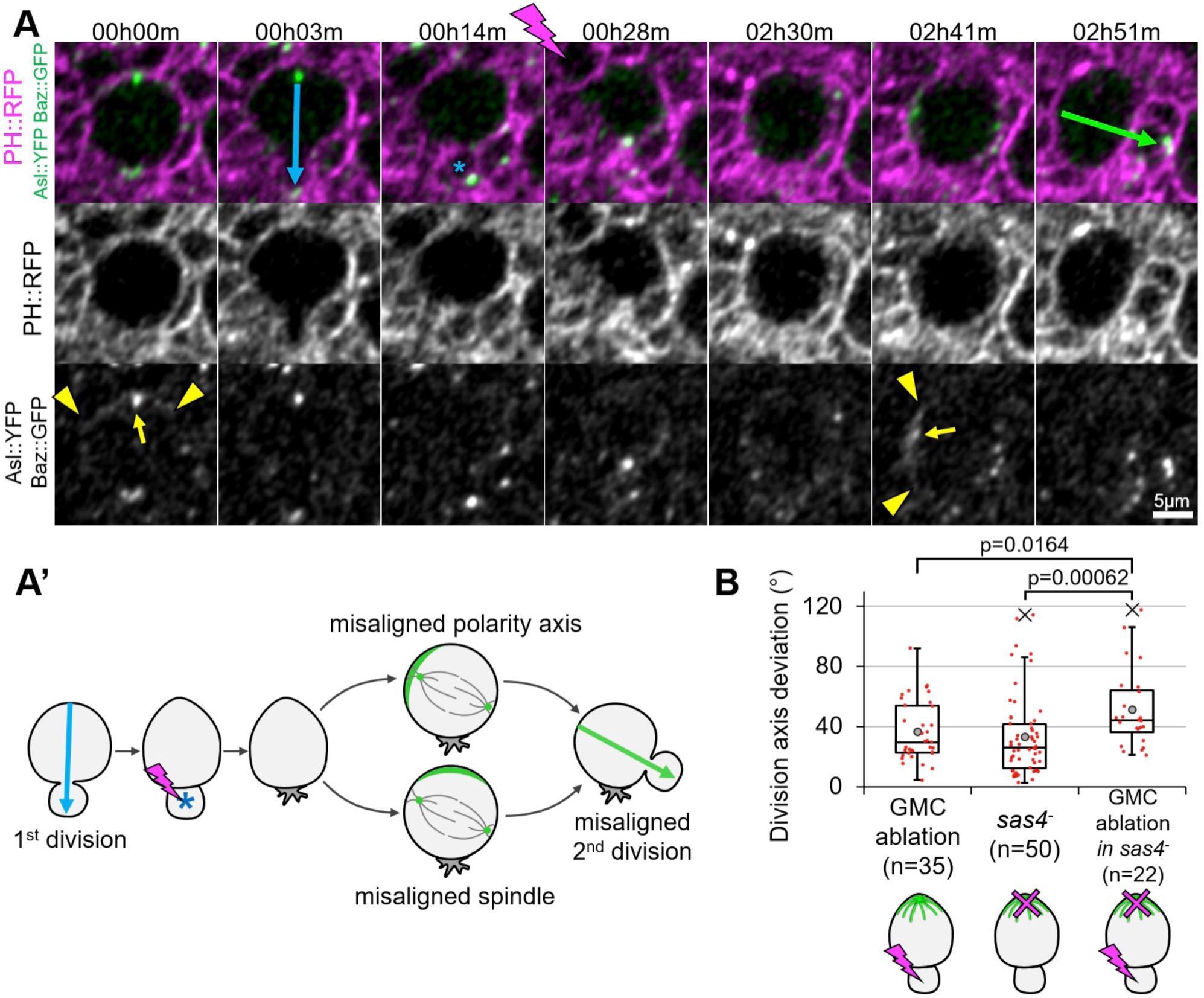
The GMC is a polarizing cue. **// A)** Two successive divisions of neuroblasts expressing the membrane marker PH::RFP (magenta), the apical marker Baz::GFP and the centrosome marker Asl::YFP (green) pre and post ablation. The ablation takes place between the two frames separated by a lightning symbol. Blue and green arrows: division axis. Yellow arrows: apical centrosome. Arrowheads: limits of Baz::GFP apical crescents. Blue asterisk: GMC prior to ablation. **A’)** Schematic of the two possibilities explaining misaligned divisions following GMC ablation. Green crescent: apical pole. Grey lines: mitotic spindle. Green dots: centrosomes. **B)** Deviation of the division axis following GMC ablation in control neuroblasts (18±14°, n=35, *also shown in* ***Figures 2A’’*** *and* ***5C***), no ablation in *sas4* mutant neuroblasts (33±26°, *n=50, also shown in* ***Figure S1B’’***) and GMC ablation in *sas4* mutant neuroblasts (49±28°, n=22). **//**

Finally, we tested whether the GMC-dependent polarizing cue acts in the same pathway as the previously reported intrinsic polarity cue for larval neuroblasts ^37^. We reasoned that, in this case, the division axis maintenance defect upon disruption of the intrinsic polarity cue division axis in *sas4* mutants should not increase upon ablation of the last-born GMC. On the contrary, the division axis deviation of *sas4* mutant neuroblasts increased significantly upon GMC ablation (**Figure 3B**). We conclude that the GMC is an extrinsic polarizing cue contributing to neuroblast division axis maintenance in parallel to the intrinsic, *sas4*-dependent cues.

### The interface of a neuroblast with its last-born daughter cell has specific features

We next sought to understand the molecular mechanism for the ability of the GMC to act as a polarizing cue. We hypothesized that the interface between the neuroblast and its latest daughter cell was likely to mediate this function, and that this interface might have specific characteristics distinguishing it from contacts between the neuroblast and older daughter cells. Certain modes of divisions in budding yeast are oriented by a “division scar” ^22^ as a landmark guiding the orientation of the next division. We hypothesized that neuroblasts could use a similar strategy, involving the midbody as a polarizing cue and, tracing the midbody marker Pavarotti-GFP ^41^, we found that the midbody was present at the newly formed neuroblast/GMC interface from cytokinesis onward and that the midbody from the previous division was inherited by the GMC (**Figure 4A**, 34/51 cell cycles). Thus, in most cases the neuroblast cortex harbours one midbody marking the position of the last-born GMC. However, tracing the fate of the midbody over several divisions in multiple neuroblasts revealed that in some cases the midbody was internalized by the neuroblast during interphase (**Figure S3A**, 17/51 cell cycles). Comparing the deviations in division orientation between neuroblasts that internalized the midbody and those that did not, revealed no significant difference (**Figure S3B**). Therefore, the midbody itself is unlikely to serve as a landmark that is directly read by the neuroblast polarizing machinery at the time of neuroblast polarization.

However, we noticed other specific features of the neuroblast/GMC interface in the vicinity of the midbody. Neuroblasts expressing membrane markers such as the GTPase Rap1 ^42^ or the PI(4,5)P2-specific PH domain of phospholipase Cδ fused to GFP (PH::GFP) (**VIDEO 6**), displayed structures resembling long tubules, largely restricted to and expanding from the neuroblast/GMC interface into the neuroblast cytoplasm (**Figure 4B**). They systematically formed around the Septin2::GFP-labelled midbody (**Figure 4C**) within the five minutes following closure of the cytokinetic furrow and were maintained throughout interphase until they disappeared when neuroblasts entered prophase (**Figure 4B**).

Further examination of the neuroblast/GMC interface revealed that it was rich in F-Actin (**Figure 4D**). This prompted us to examine the subcellular localization of several actin regulators using endogenously expressed fluorescent protein traps. We found Flare (actin depolymerizing factor, ^43^), Canoe (Afadin, ^44^) and Cindr ^45^ (**VIDEO 7**) were present at the neuroblast/GMC interface (**Figure 4D**).

We further reasoned that adhesion molecules could be involved in division orientation maintenance and might be enriched at the neuroblast/GMC interface. We focused on the adhesion molecule E-Cad, as it is involved in orienting mitosis in the fly sensory organ precursor lineage ^16^, is expressed in neuroblasts and their lineage and has been reported to be enriched between neuroblasts and their daughter cells ^46,47^. However, neither E-Cad nor its binding partner β-Catenin were restricted to the interface of neuroblasts and the last-born GMC compared to interfaces between neuroblasts and older GMCs (**Figure S4A-B**). In conclusion, specific characteristics distinguish the interface between the neuroblast and its last-born daughter cell: the presence of a midbody, plasma membrane extensions, and an accumulation of actin and actin-regulators.

### Components of the neuroblast/GMC interface polarize neuroblasts

We next used neuroblast and neuroblast progeny-specific RNAi to test whether components of the midbody and the neuroblast/GMC interface are involved in orienting the axis of neuroblast divisions. Efficient depletion of E-Cad by RNAi (**Figure S4B**) did not result in significant division orientation defects of central brain neuroblasts (**Figure S4C**). Therefore, despite being involved in controlling niche position of mushroom body neuroblasts ^48^, E-Cad is not critical for neuroblast division orientation maintenance in the central larval brain.

Surprisingly, although midbody internalization had no effect on division axis maintenance (**Figure S3**), efficient depletion of the midbody components Septin 1 (**Figure S4DD’**) as well as RNAi against Septin 2 led to significant division orientation defects (**Figure 5A**). Septins are required for cytokinesis in certain tissues ^49^, raising the possibility that cytokinesis failure indirectly disrupts division axis orientation. However, we always observed complete and stable furrow ingression (not shown) and never observed multinucleation in Septin1 and Septin2-depleted neuroblasts (**Figure S4**). This is either because depletion of Septins was only partial (**Figure S4D-D’**) or because Septins are not required for cytokinesis in neuroblasts, as it is the case in other contexts ^50-52^. Furthermore, efficient depletion of Flare and Cindr (**Figure S4EF**) resulted in a significant increase of the division axis deviations (**Figure 5A**). This prompted to express RNAi against a Cindr interactor, the transmembrane immunoglobulin Roughest (Rst, ^53^), and against its adaptor Dreadlock (Dock, ^54^), depletion of all of which also affected division axis maintenance (**Figure 5A**).

Like misaligned divisions following ablation of the GMC, misaligned divisions caused by RNAi against Flare, Cindr, Rst and Dock were associated with misplaced apical crescents rather than spindle-cortical polarity alignment problems (**Figure 5B, VIDEO 8**). We further reasoned that misplaced apical crescents, instead of being caused by the disruption of an additional polarizing cue relying on the neuroblast/GMC interface, could be caused by an abnormally mobile apical centrosome directing polarization at the wrong place (**Figure S4G**). However, although we did occasionally observe apical centrosome detachment during interphase, this occurred at the same frequency in control and RNAi expressing neuroblasts (control: 4/68; Sep1 RNAi: 4/47; Sep2 RNAi: 2/53; Cindr RNAi: 6/86; Dock RNAi: 7/73; Rst RNAi: 2/53; Flare RNAi: 6/55). Furthermore, precise measurements of the apical centrosome position suggest that, at least in Flare-depleted neuroblasts, the apical centrosome position when neuroblasts polarize does not correspond to the apical crescent position in metaphase (**Figure S4G**). Therefore, misplaced Baz crescents upon RNAi depletion of these components are unlikely to be caused by an incorrectly positioned apical centrosome.

Finally, we hypothesized that, if depleting components of the neuroblast/GMC interface disrupts the role of the GMC as a polarizing cue, GMC ablation together with depleting these components should not further increase the resulting deviations of the orientation of neuroblast division. Consistent with this idea, GMC ablation in Sep1-depleted neuroblasts did not significantly increase division axis deviations when compared to Sep1-depleted neuroblast alone or control neuroblasts on which GMC ablations were performed (**Figure 5C**). Based on these results, we propose that the role of the GMC in maintaining the division axis of neuroblasts is mediated by proteins specifically localizing to the newly formed interface between the neuroblast and its last-born daughter cell (**Figure 5D**).

## Discussion

Deciphering the signals that provide positional information is a central issue in understanding how cell divisions are oriented. Here, we addressed this question in the highly proliferative neuroblast in the *Drosophila* larval brain. Our results provide an explanation for why maintenance of the orientation of divisions is only partial disrupted in larval neuroblasts lacking the intrinsic polarizing cues ^37^: the last-born daughter cell of neuroblasts acts as an additional, extrinsic polarizing cue (**Figures 2,3**). Consistent with this, disruption of both the intrinsic and extrinsic polarizing cues increases the defects of division axis maintenance when compared to disrupting each cue alone (**Figure 3B**). It is noteworthy, however, that this does not result in complete randomization of division orientation, suggesting that either polarizing cues are not fully disrupted and/or that additional polarizing cues contribute to the process.

We find that larval neuroblasts are not exclusively polarized by cell intrinsic mechanisms, since an external signal received at the basal pole biases the decision of which part of the neuroblast cortex will become apical in the following mitosis. The exact molecular nature of the polarizing cue is unclear. Although bearing similarities with division axis maintenance in budding yeasts, relying on a Septin-rich cytokinesis remnant ^22^, the midbody of neuroblasts is unlikely to be the landmark. Indeed, midbody internalization does not affect division axis maintenance (**Figure S3**), and depletion of Cindr or Flare, which do not localize to the midbody, affects maintenance (**Figure 5A**). Therefore, the cue may rely on other components initially assembled by the midbody. This could be cell-cell contacts, explaining the involvement of an adhesion molecule such as Roughest (**Figure 5A**). This would further explain why GMC ablation, although not directly targeting the interface, affects division axis maintenance (**Figure 2A,B**), and is consistent with the role of the midbody in organizing cell-cell contacts ^55^. It also seems likely that the midbody organizes the actin-rich plasma membrane extensions, given the physical origin (the midbody) and timing (immediately after cytokinesis) of their appearance (Figure 4, **VIDEO 6**). Interestingly, Septins were shown to physically interact with the mammalian orthologs of Cindr and Roughest ^56-58^, providing a possible link between the Septin-rich midbody and the Cindr-rich neuroblast/GMC interface.

**Figure 4:**
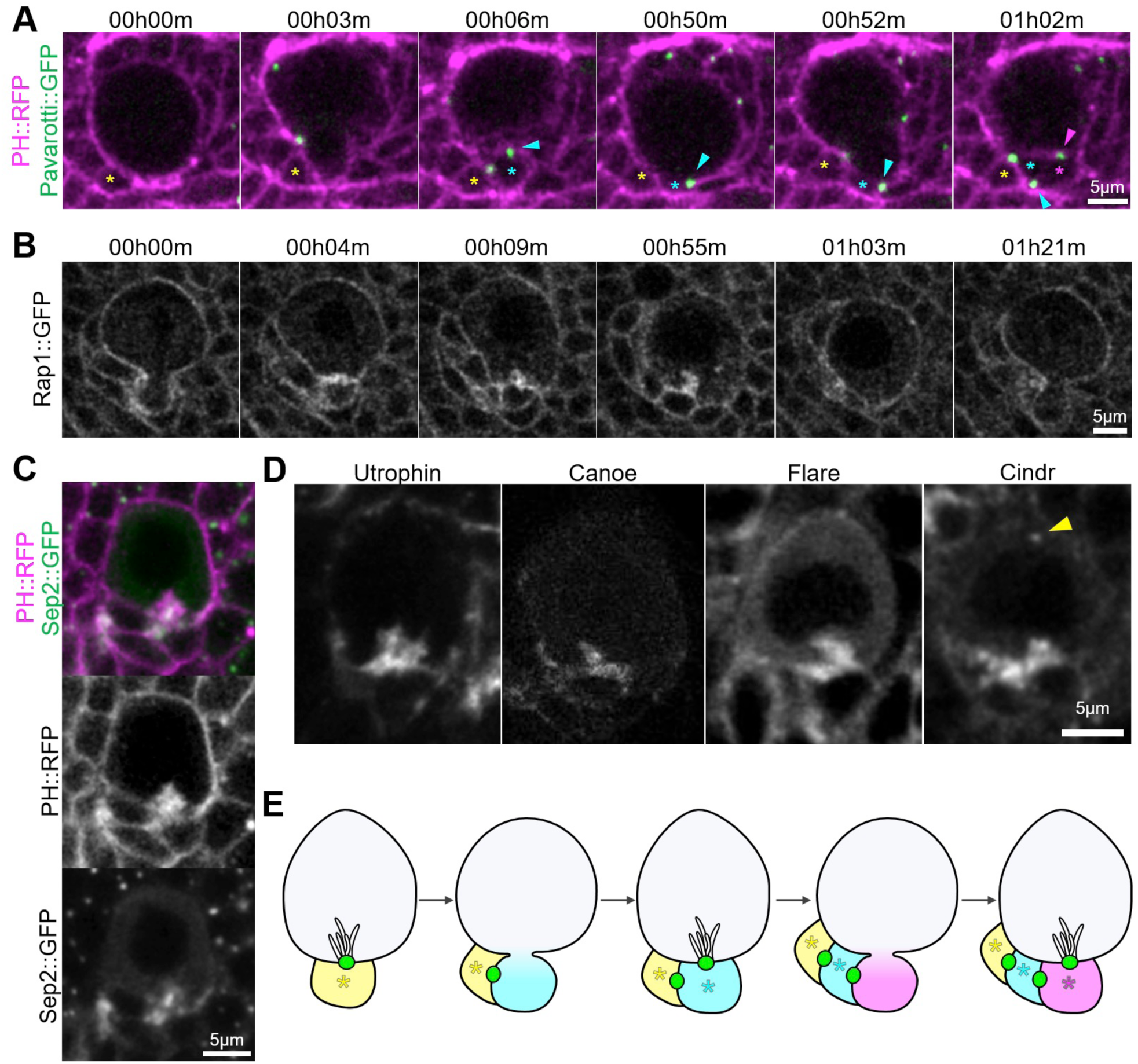
The neuroblast/GMC interface presents specific characteristics. **// A**) Two successive divisions of neuroblasts expressing the membrane marker PH::RFP (magenta) and the midbody marker Pavarotti::GFP (green). Yellow asterisk: previously born GMC, cyan asterisk: GMC formed at subsequent division, magenta asterisks: last-born GMC in this series. Blue arrowhead: midbody formed at the first division recorded stays at the newly formed neuroblast/GMC interface, but is inherited by the GMC in the next division, when a new midbody remains at the subsequently formed neuroblast/GMC interphase (magenta arrowhead). **B)** Neuroblast expressing the membrane marker Rap1::GFP. Tubular plasma membrane extensions originating from the neuroblast/GMC interface appear within ˜ 5 minutes following closure of the cytokinesis ring (00h04m-00h09m), persist throughout interphase (00h55m) and disappear upon cell rounding at the onset of mitosis (01h03m). **C)** Interphase neuroblast expressing Sep2::GFP (green) and the membrane marker PH::RFP (magenta). **D)** Interphase neuroblasts expressing GFP or YFP-fusions to the indicated proteins localizing to similar tubular extensions at the neuroblast/last-born GMC interface. Cindr::GFP also localizes to centrosomes (arrowhead). **E)** Schematic illustrating the specific characteristics of the neuroblast/last-born GMC interface and the inheritance of this interface by the GMC at each division. Green: midbody. Asterisk: GMCs, same color code as figure 3A. **//**

How can a basally received cue direct apical polarization, at the opposite pole of the neuroblast? This is reminiscent of the polarization of the *C. elegans* zygote: the sperm entry point acts as a cue inducing an actomyosin flow ^20^ that establishes Par complex polarity at the opposite end of the cell (^59^ for review). Interestingly, Septins control some parameters of this flow ^60^. This “long-range” control of polarity further involves tensions in *C. elegans* ^61,62^. In migrating neutrophils, polarity maintenance also relies on the propagation of mechanical force from one pole to the opposite one ^63^. Future work will explore whether Actomyosin/Septin-dependent flows and/or propagating forces similarly mediate polarization of neuroblast.

Finally, is division axis maintenance in larval neuroblasts physiologically important? In the brain, neuroblasts and their progeny are ensheathed by glial cells ^64^ that provide proliferative signals ^65^ protection against starvation ^66^ and oxidative stress ^67^. By placing daughter cells between the neuroblast and their glial cell niche, randomized division orientation could isolate the neuroblasts from the protective influence of glial cells. We further speculate that, contrary to other stem cells using contact with their niche as a spatial cue ^19^ to orient their division, the surface contact of the neuroblasts/ glial niche is too large to provide an accurate spatial cue. In this situation, reading the orientation of previous divisions using the contact with the last-born daughter cell as positional landmark appears as an efficient strategy to maintain stem cells within range of those signals. Therefore, it would be of particular interest to explore in other stem cell models whether similar strategies are used.

## Methods

### Fly stocks and genetics

Flies were reared on standard corn meal food at 25°C, except for RNAi-expressing larvae and their corresponding controls (Figures 5, **S4**), which were placed at 30°C from the L1 larval stage to the L3 stage, at which point they were dissected. As RNAi was driven using the Worniu-GAL4 driver, which is not expressed in every neuroblasts, UAS-nls::BFP was used as a GAL4 reporter to identify and exclude from the analysis neuroblasts not expressing GAL4.

**Figure 5:**
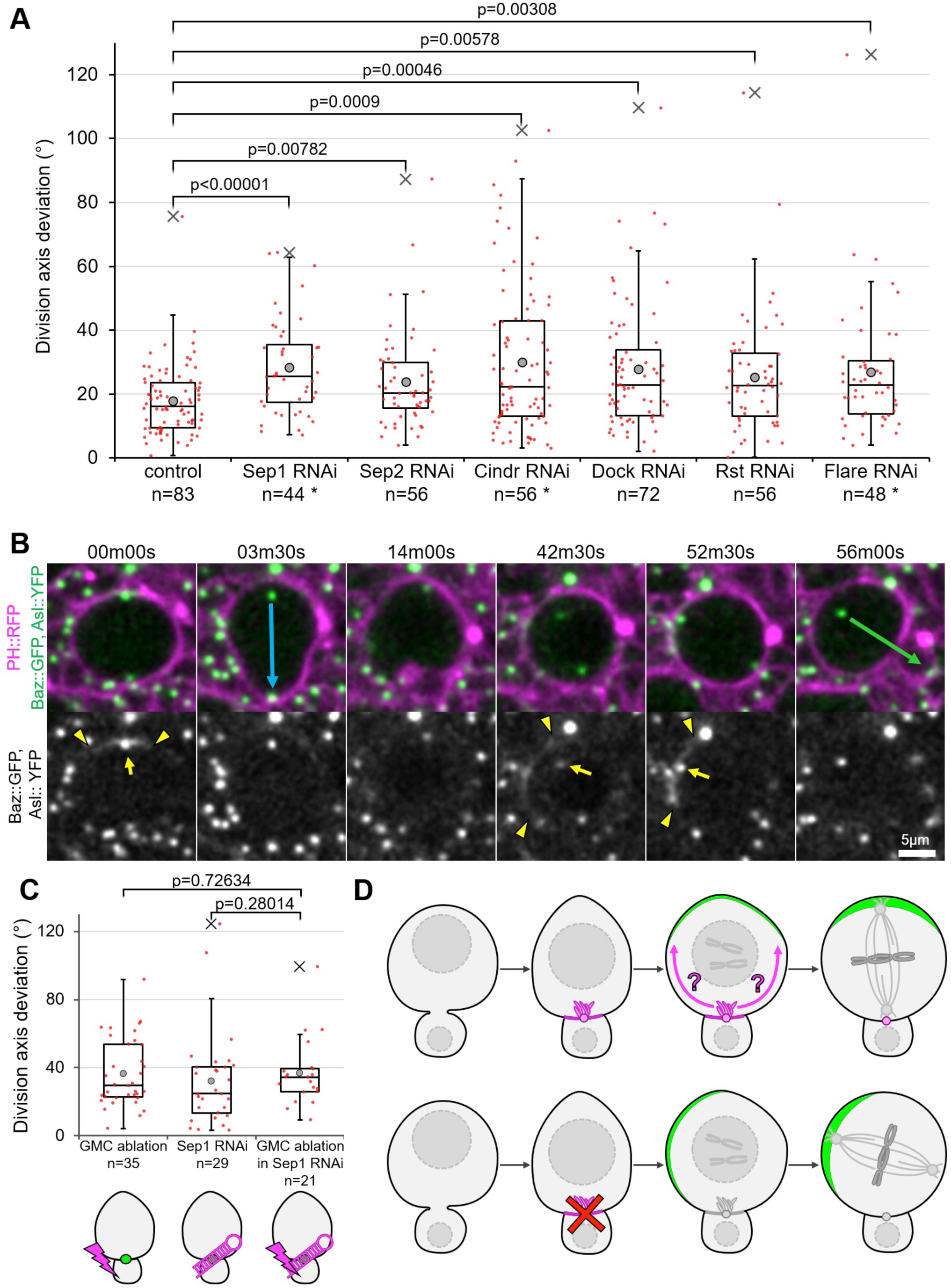
Components of the neuroblast/GMC interface polarize neuroblasts. **// A)** Deviation of the division axis from one cell cycle to the next in neuroblasts expressing *worniu*-Gal4-driven RNAi against the indicated components of the neuroblasts/GMC interface, as well as the Cindr interactors Rst and Dock. Asterisks: candidates for which protein depletion was confirmed. See materials and methods for precise genotypes. **B**) Two successive division of a neuroblast expressing *worniu*-Gal4-driven Cindr RNAi. The second division axis (green arrow) deviates from the first one (blue arrow) by 53°. The position of the apical Baz crescent (delimited by arrowheads) has shifted, whereas the apical centrosome (arrow) aligns properly with it. Correct alignment of cortical polarity and the mitotic spindle was observed in all RNAi knockdown experiments shown in A (not shown). **C**) Deviation of the division axis following GMC ablation in control neuroblasts (36±20°, n=35, *also shown in* ***Figures 2A’’*** *and* ***3B***), in neuroblasts expressing *worniu*-Gal4-driven Sep1 RNAi (32±27°, n=29) and following GMC ablation in neuroblasts expressing *worniu*-Gal4-driven Sep1 RNAi (37±19°, n=21). **D)** Graphical summary of findings. Top row: the neuroblast/last-born GMC interface, distinguishable from other interfaces by the presence of the midbody and actin-rich plasma membrane extensions (magenta), biases through an unknownmechanism (magenta arrows) the apical polarization (green crescent) of neuroblasts as they round up in mitosis. The mitotic spindle aligns with this crescent, resulting in division axis maintenance. Bottom row: disruption of this mechanism leads to misplaced apical polarization (with which the mitotic spindle still properly aligns) resulting in defective division axis maintenance. **//**

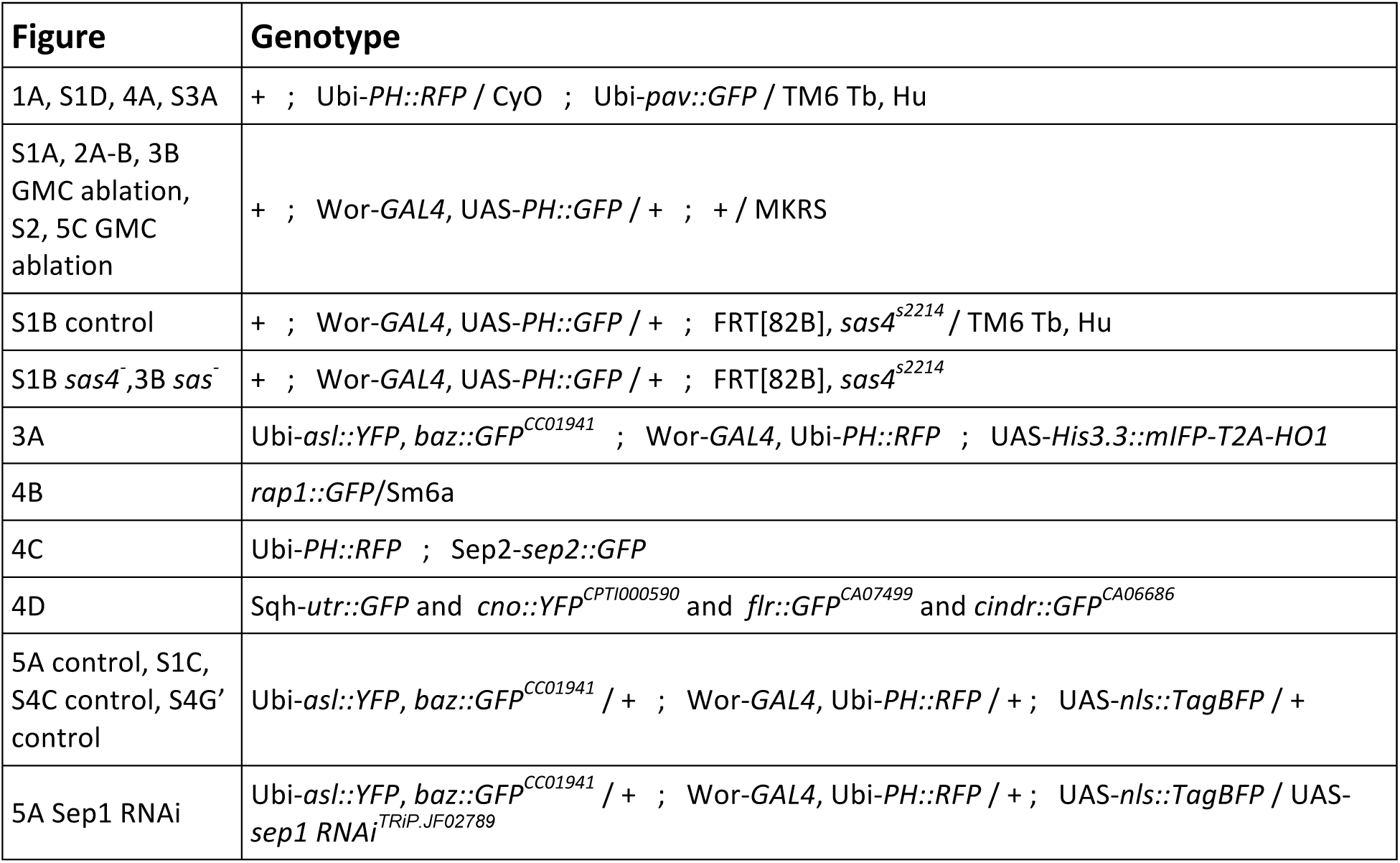

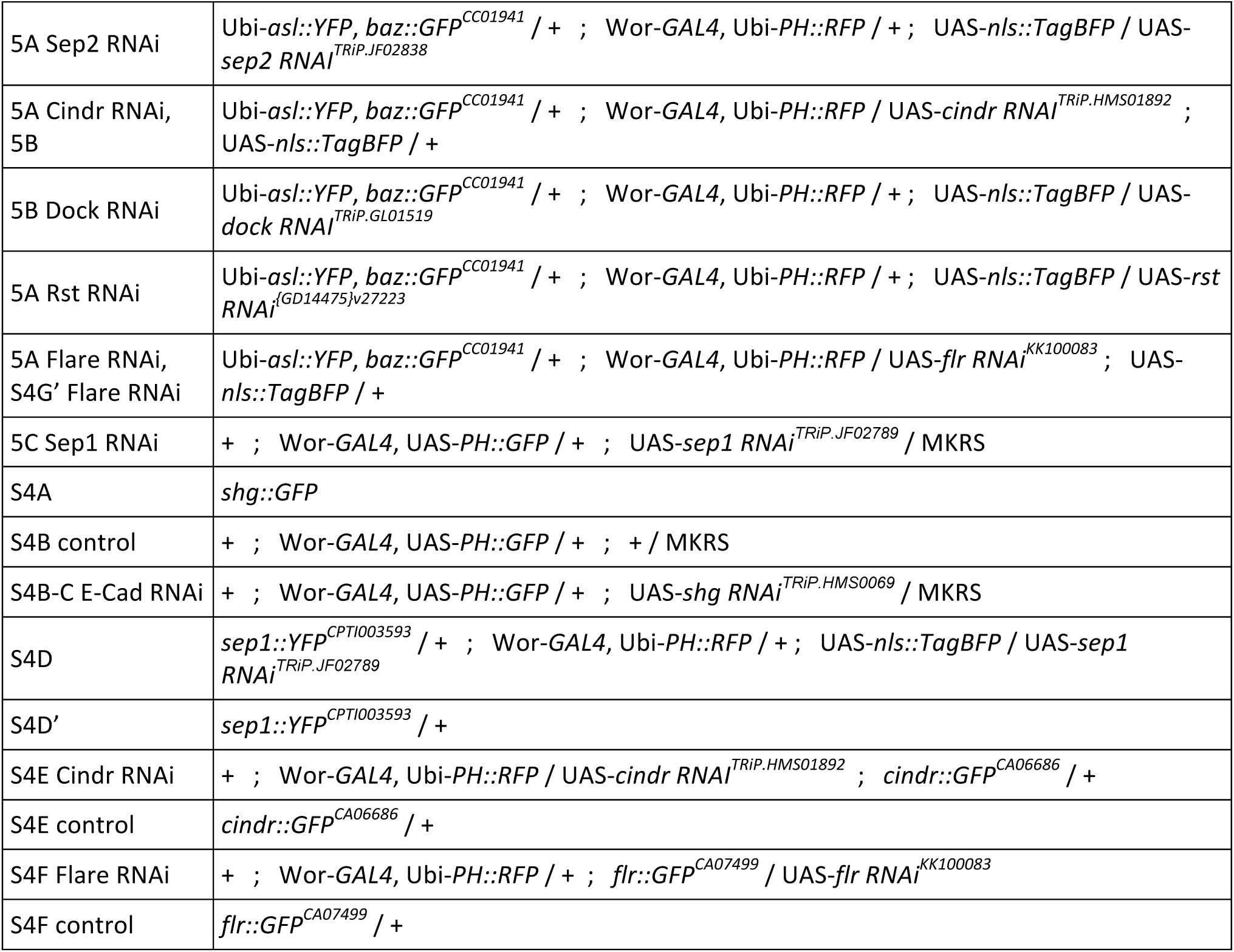

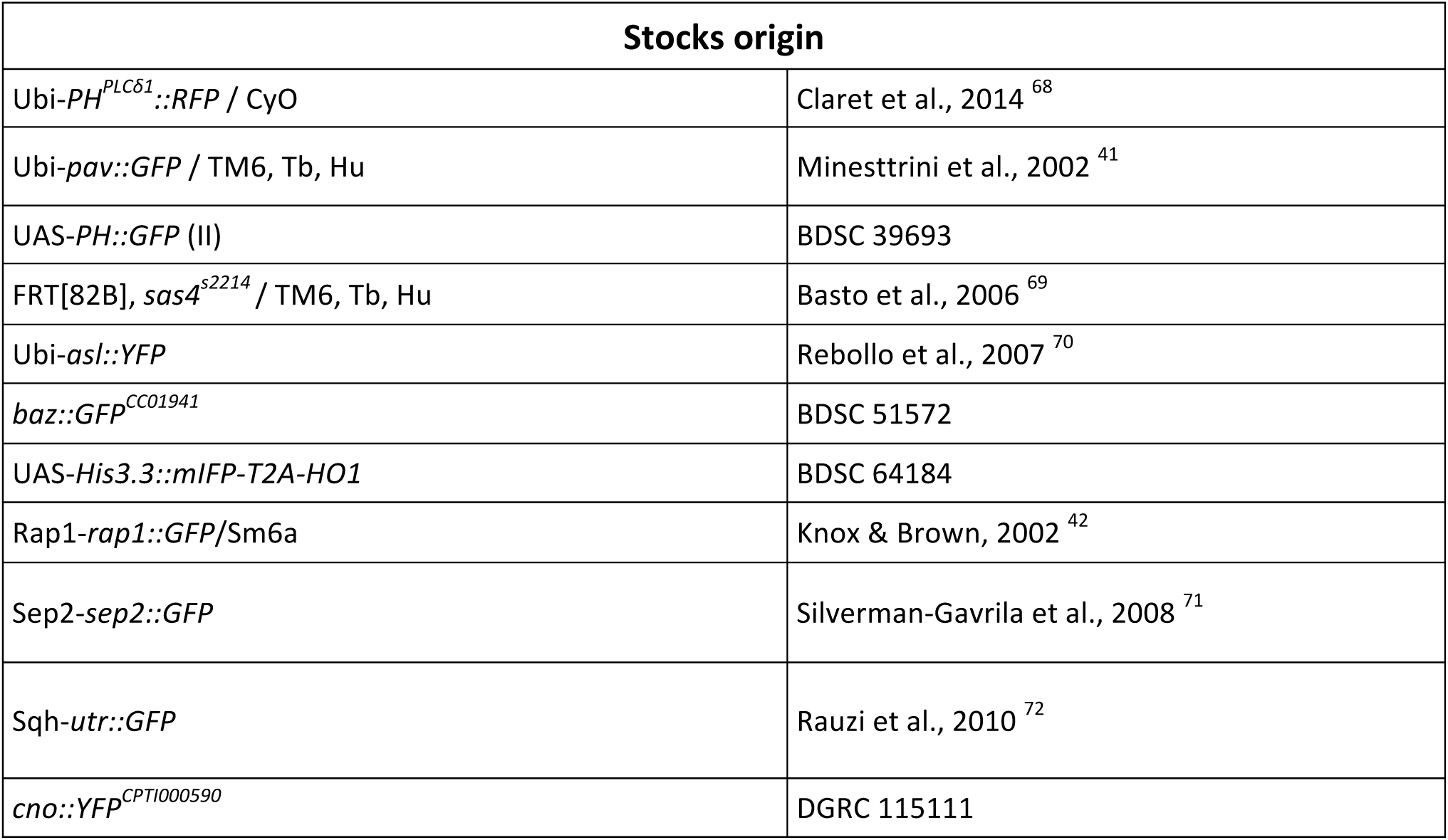

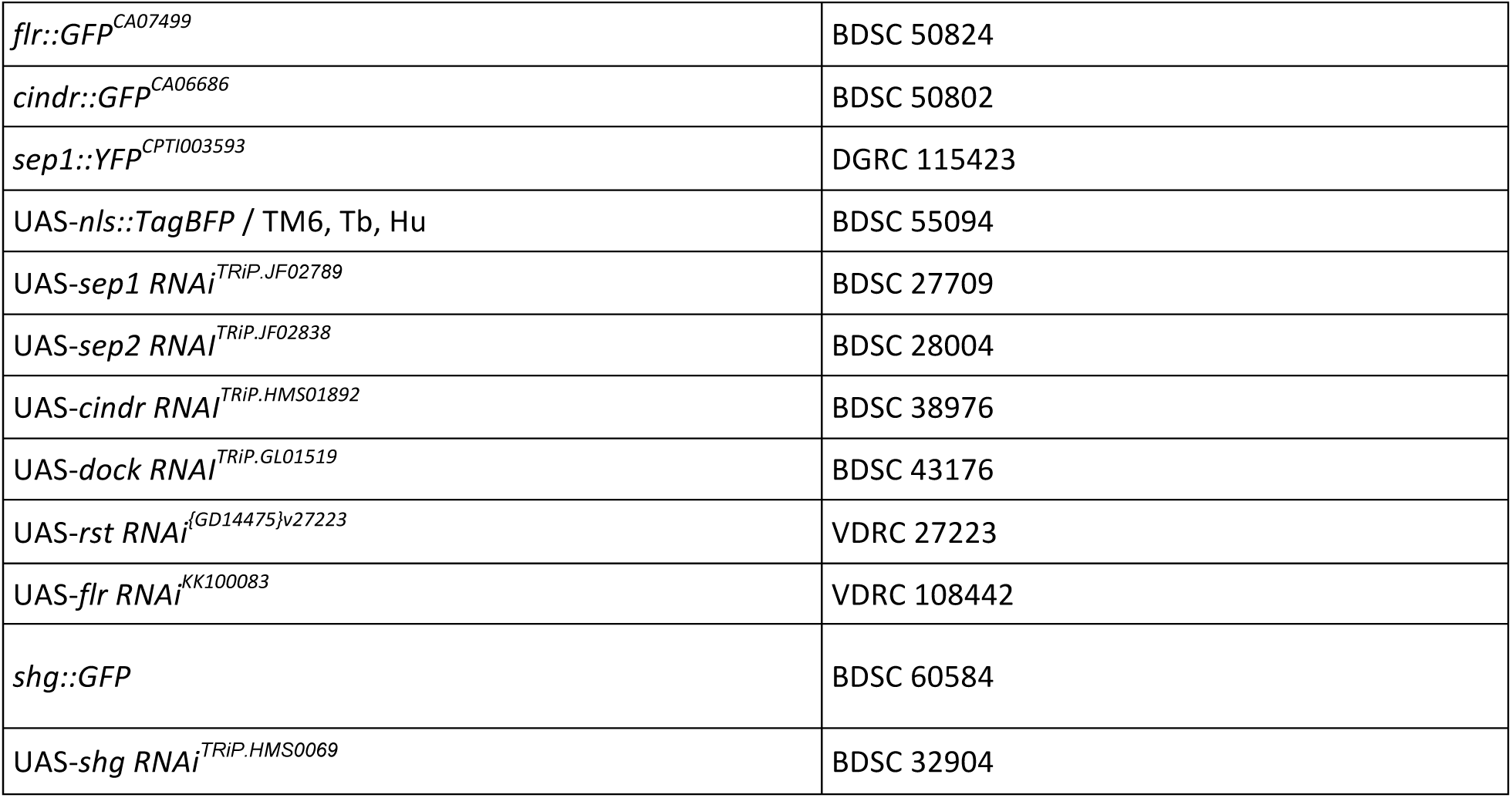

### Live imaging

Every reference to Schneider’s medium corresponds to glucose-supplemented (1g.l^-1^) Schneider’s medium (SLS-04-351Q). Live imaging was performed as described ^73^. Entire brains were dissected from early L3 larvae (still crawling inside the food) in Schneider’s medium, isolated from the surrounding imaginal discs. Particular care was taken to avoid pulling on brains at any time during the dissection and damaged brains were discarded. Isolated brains were transferred to a drop of fibrinogen dissolved in Schneider’s medium (50mg.ml^-1^) on a 25mm Glass bottom dish (WPI), which was then clotted by addition of thrombin (100Uml-1, Sigma T7513). Clots were then covered in Schneider’s medium (approximately 750µl spread over the entire surface of the glass).

RNAi-expressing brains and their associated controls (**Figures 5, S4**) were then imaged on a LEICA SP8 confocal microscope (LEICA) equipped with a 63x NA 1.2 water immersion objective lens. Stacks of 25-30 optical z-section separated by 0.8µm, covering a 132×132x20-24µm region of the surface of the antero-ventral central brain were acquired every 210s for 2h30m to image Asl::YFP, Baz::GFP and PH::RFP, after which a final stack also imaged the GAL4 reporter Nls::BFP. For laser ablations (**Figures 2, 3, 5C, S2**), brains were imaged on a Zeiss 710 confocal microscope equipped with a 63x oil immersion objective lens. Stacks of 16 optical z-section separated by 1.2µm, covering a 75×75×18µm region of the surface of the antero-ventral central brain were acquired every 210s for 2h30m to 3h.

### Laser ablation

Laser ablations were performed on a 710 confocal microscope (Zeiss) equipped with a 63x oil immersion objective lens and a two-photon tunable Chameleon from Coherent set to 800nm, using the fluorescence recovery after photobleaching (FRAP) module of the Zen software. Settings were as follow: laser intensity 25-32% (empirically adjusted depending on the depth of the targeted area within the tissue; targeted area 1.4×1.4µm; 15 iterations.

### Image processing and angle measurement

Data was processed and analyzed using ImageJ ^74^. A 0.8×0.8×0.8 pixels wide 3D gaussian blur was applied to every image. The 3D vectors corresponding to the division axis were defined by the 3D coordinates of the apical and the basal pole at telophase when only a membrane marker was available (see **Figure S1A**), or of the apical and basal centrosome at metaphase when a centrosome marker was available. The angle (α) between two 3D vectors was calculated using the formula:

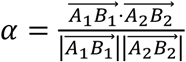

where the dot product is:

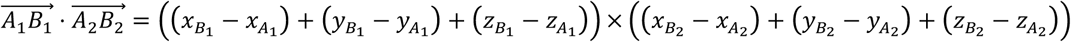

and the magnitude of any vector is:

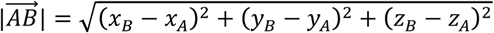

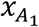 being for example the x coordinate of the apical centrosome during the first division, and 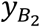 being for example the y coordinate of the basal centrosome during the second division.

### Statistical analysis

P values displayed over boxplots were calculated using a non-parametric two-tailed Mann-Whitney U Test. The numbers displayed in boxplots correspond to measurements in individual neuroblasts. The number of animals used in each experimental dataset is as follows: **Fig 1**: 2. **Fig 2A-A’’** no ablation: 10; control ablation: 7; GMC ablation: 11. **Fig 2B-B’’**: 4. **Fig 3A**: 2. **Fig3B** GMC ablation: 11; *sas4*^*-*^: 7; GMC ablation in *sas4*^*-*^: 8. **Fig. 4A**: 2. **Fig. 4B**: 3. **Fig. 4C**: 2. **Fig. 4D** Utrophin: 2; Canoe: 5; Flare: 6; Cindr: 6. **Fig 5A** control: 4; Sep1 RNAi: 4; Sep2 RNAi: 2; Cindr RNAi: 3; Dock RNAi: 3; Rst RNAi: 2; Flare RNAi: 3. **Fig 5B**: 3. **Fig 5C** GMC ablation: 11; Sep1 RNAi: 4; GMC ablation in Sep1 RNAi: 4. **Fig. S1B-B’’**: 4. **Fig. S1B-B’’** control: 10; *sas4*^*-*^: 7. **Fig S1C**: 3. **Fig S1D**: 2. **Fig S2** GMC ablation: 10; control ablation: 7. **Fig. S3A-B**: 2. **Fig S4A**: 2. **Fig S4B** control: 2; E-Cad RNAi: 2. **Fig S4C** control: 4; E-Cad RNAi: 3. **Fig S4D**: 2. **Fig S4D’**: 1. **Fig S4E** Cindr RNAi: 1; control: 1. **Fig. S4F** Flare RNAi: 2; control: 3. **Fig S4G** control: 4; Flare RNAi: 3.

### Immunostainings

For the β-Cat staining (**Figure S3B**), brains were dissected in PBS, fixed for 20 minutes in 4% formaldehyde (Sigma), permeabilized in PBS-Triton 0.1% (PBT) for 1 hour, incubated in a Rabbit-anti-β-Cat^central^ antibody ^75^ diluted 1:200 in PBT for 2 hours, rinsed 3 times in PBT, washed 3 times in PBT for 10 minutes, incubated in a secondary Donkey-anti-Rabbit antibody coupled to Alexa 594 (Thermo Fisher) diluted 1:1000 in PBT for 1 hour, rinsed 3 times in PBS, washed 3 times in PBS for 10 minutes, transferred to a 50% glycerol solution, and mounted in a Vectashield mounting medium (Vector Laboratories). Every step was performed at room temperature.

## Acknowledgements

We would like to thank A.Guichet, N.Brown, J.Raff, C.Doe, C.Gonzalez, T.Lecuit, A.Wilde and D. Glover for providing reagents and I.Näthke for critical reading and K.Mckay for help with the RNAi screen. We also thank the Bloomington stock center and the VDRC for providing fly lines and P.Appleton and S.Swift from the Center for advanced Scientific Technologies for technical support for microscopy. Work in JJ’s lab is supported by a Sir Henry Dale fellowship (100031Z/12/Z). The tissue imaging facility is supported by the grant WT101468 from Wellcome.

## Author contribution

N.L and J.J. designed the study. N.L designed and performed the individual experiments. N.L. and J.J. analysed and interpreted the data and wrote the manuscript. J.J. acquired funding.

## Supplementary figure legends

**Figure S1:**
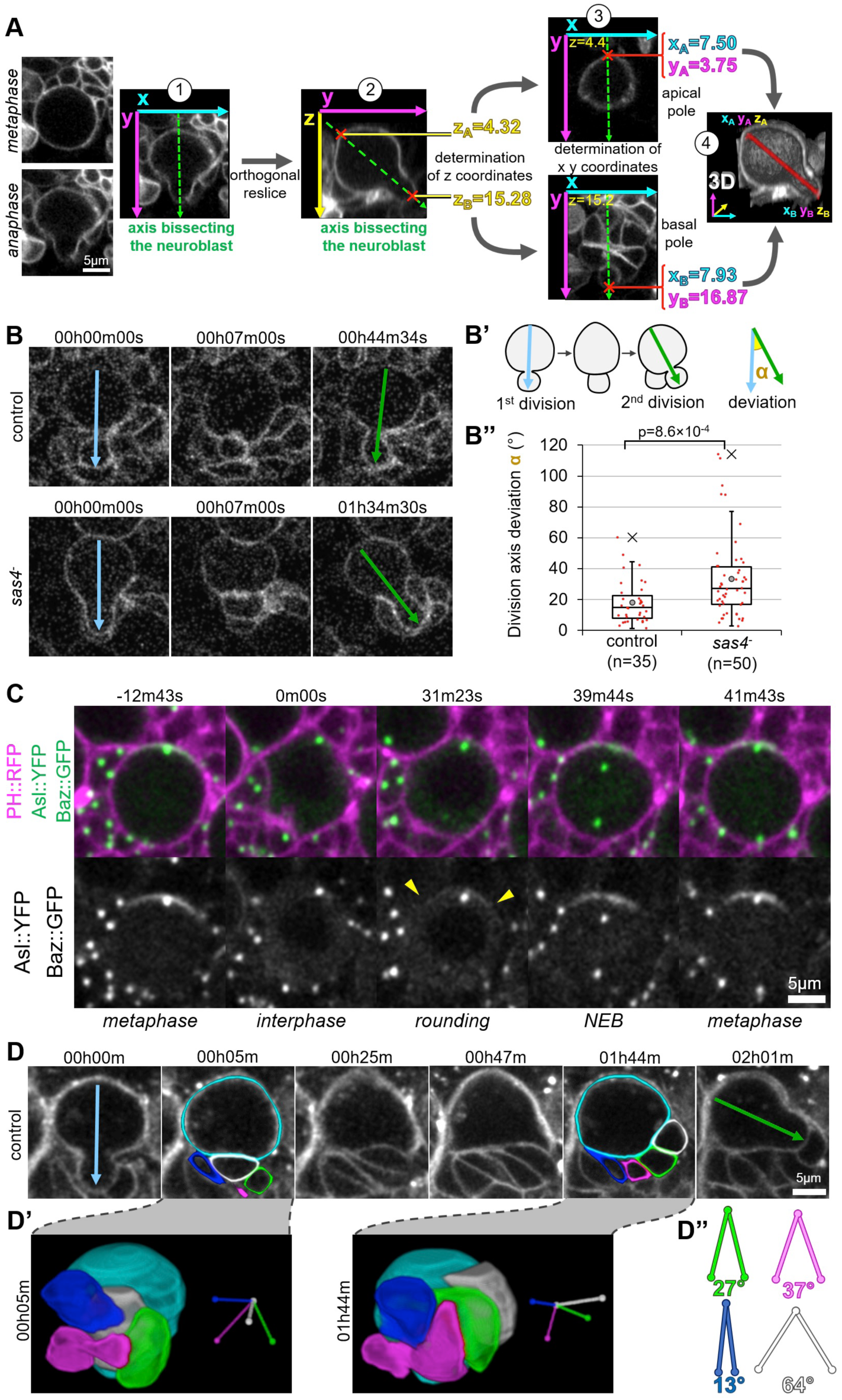
3D division angle measurements in larval neuroblasts. A) Methodology for 3D division axis measurement. 1) At telophase, a manually determined axis bisecting the neuroblast (green arrow) is used to orthogonally slice a 3D stack covering its entire volume. 2) This orthogonal slice is used to determine the z coordinates of the apical and basal poles (red crosses). 3) These z coordinates are used on the corresponding slices of the 3D stack to determine the x and y coordinates of the basal and apical poles (red crosses). 4) 3D reconstruction of the neuroblast volume and the division axis measured with this method (red). Angles between successive division axis are calculated from these 3D coordinates (see Methods). **B)** Two successive divisions of control and *sas4* mutant neuroblasts expressing *worniu*-GAL4-driven PH::GFP. **B’**) Schematic of the angles α measured in the movies shown in the previous panels. **B’’)** Distribution of the angles α described in the previous panels in control (18±14°, *n=35, also shown in* ***Figure 2A’’***) and *sas4* mutant neuroblasts (33±26°, n=50, *also shown in* ***Figure 3B***). **C)** High time resolution movie (Δt: 23.84s) of a neuroblast expressing PH::RFP (magenta), Asl::YFP and Baz::GFP (green in merge). 0m00s corresponds to the beginning of interphase, when the neuroblast adopted a stable shape. A faint apical Baz crescent (delimited by arrowheads) is detected as soon as neuroblasts start rounding up. **D)** Manual segmentation of the neuroblast/progeny cluster shown in **Figure 1A** at the beginning of interphase (5 minutes after the first division) and the beginning of the second division, detected by mitotic rounding (1h44m after the first division). Cyan: neuroblast. White: last-born GMC. Magenta, green, blue: other neuroblast-neighboring cells. **D’)** 3D reconstruction of the manual segmentation shown in the previous panel (left) and 3D vectors defined by the coordinates of the barycenter of the neuroblast (where all vectors converge) and other neighboring cells (right). **D’’)** Angles between the neuroblast-other cell vectors at the two time points described in the previous panel. The GMC (white) shows a higher angle than other cells, consistent with its apparent relocalisation.

**Figure S2:**
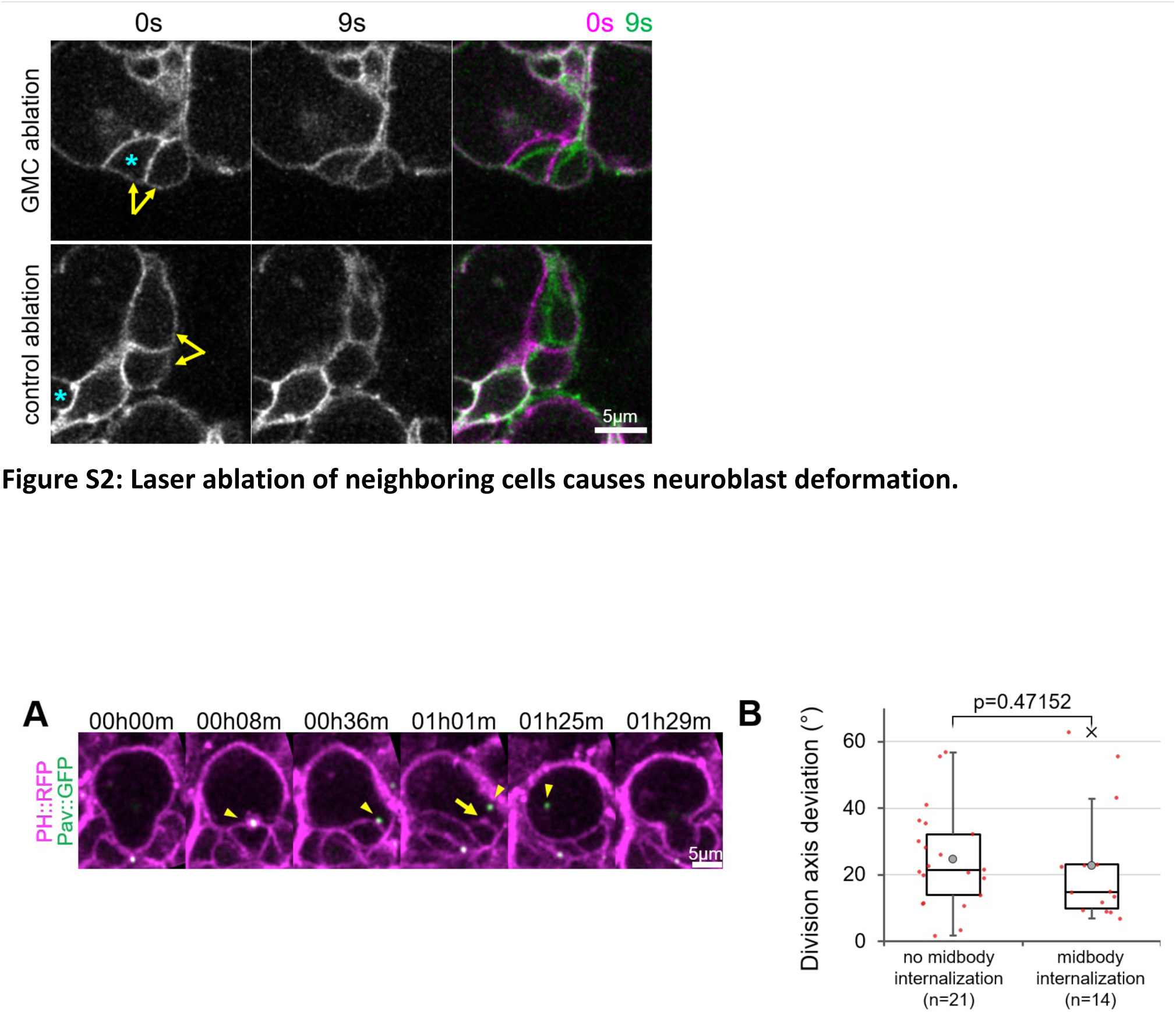
Laser ablation of neighboring cells causes neuroblast deformation. Deformation of neuroblasts expressing *worniu*-GAL4-driven PH::GFP following laser ablation, which occurs approximately 1.5 seconds after the first time point following ablation (0s). For efficient and reproducible ablations, we always targeted the interface between two cells (arrows) of the neuroblast/progeny cluster: control ablations targeted two neuroblast-neighboring cells away from the GMC, and GMC ablations targeted the GMC and another neuroblast-neighboring cell. The third panel is a composite picture of the first two panels, illustrating the immediate neuroblast deformation toward the targeted cells. Blue asterisk: last-born GMC. Yellow arrows point at relevant cells.

**Figure S3:**
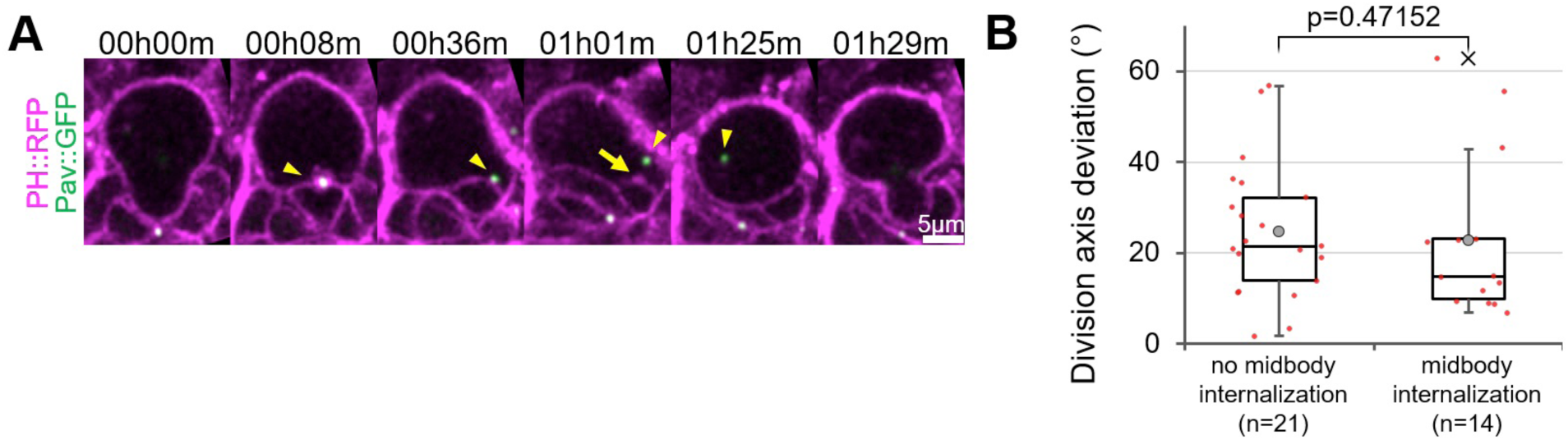
Midbody internalization does not affect neuroblast division axis maintenance. **A)** Two successive divisions of a neuroblast expressing the membrane marker PH::RFP (magenta) and the midbody marker Pav::GFP (green). The midbody formed during the first division detaches (arrowhead) from the neuroblast/GMC interface before the second division. Plasma membrane extensions (arrow) are still present after midbody internalization. **B)** Division axis deviation when the midbody is kept at the interface until the next division (25±15°, n=21) and when the midbody is internalized before the next division (23±17, n=14).

**Figure S4:**
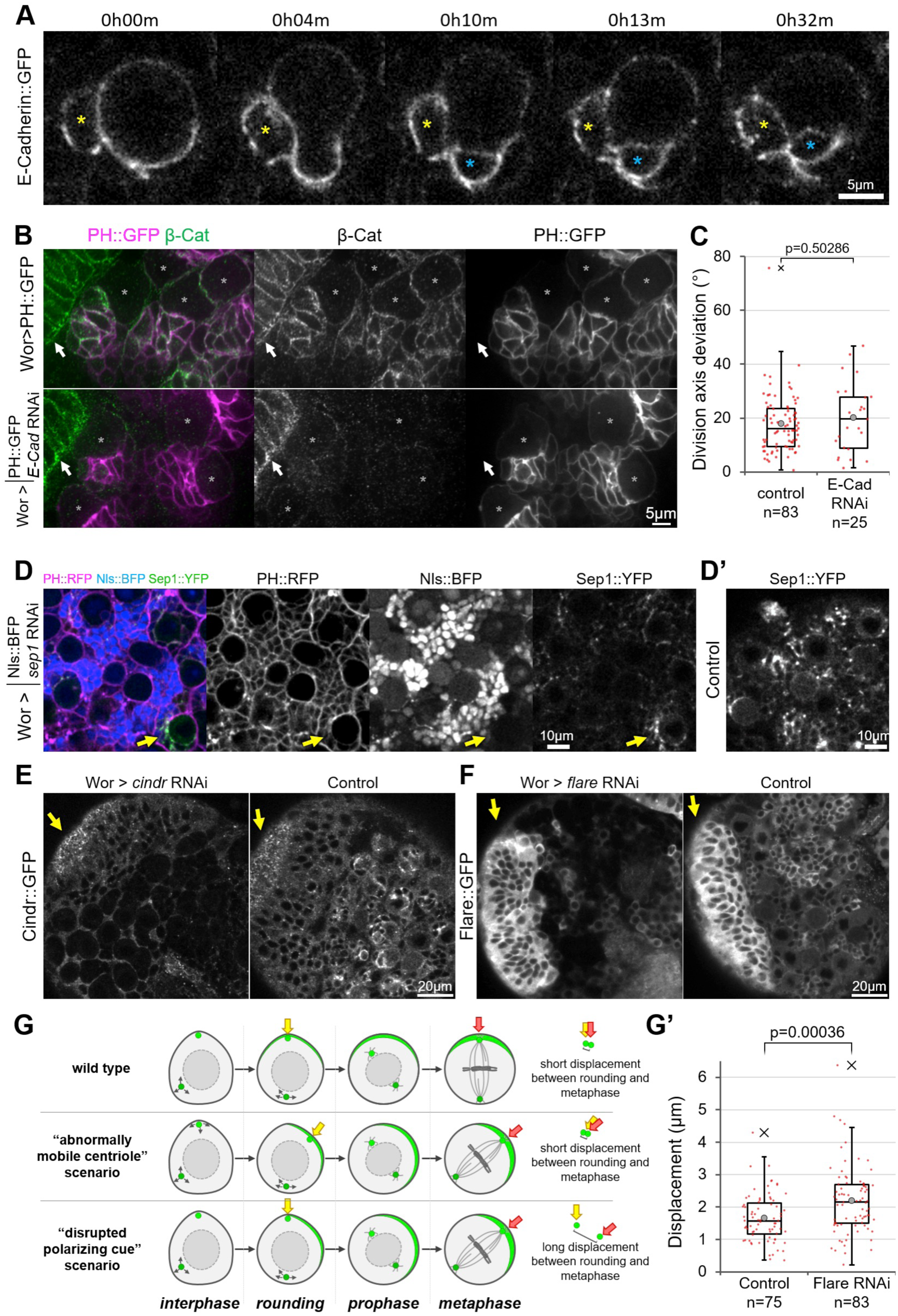
RNAi-mediated depletion of components of the neuroblast/ GMC interface. **A**) Neuroblast expressing E-Cadherin::GFP. Yellow asterisk: last-born GMC. Blue asterisk: older GMC. **B)** Neuroblasts stained for β-Cat and expressing *worniu*-Gal4-driven PH::GFP and, for the bottom panel, *worniu*-Gal4-driven *E-Cadherin* RNAi. Arrows point to a neuroepithelial region of the brain, expressing β-Catenin::YFP but not *worniu-*GAL4. **C)** Deviation of the division angle in control (18±11°, n=83, *also shown in* ***Figure 5A***) and E-Cad-depleted neuroblasts (20±13°, n=25). **D)** Neuroblasts expressing PH::RFP, Sep1::YFP, and *worniu*-Gal4-driven Nls::BFP and *sep1* RNAi1. The arrow points to a neuroblast expressing no detectable amount of the GAL4 activity reporter UAS-Nls::BFP and displaying higher amounts of Sep1::YFP. **D’)** Control neuroblasts expressing Sep1::YFP. **E)** Control and *cindr* RNAi-expressing neuroblasts expressing Cindr::GFP. The arrows point to a neuroepithelial region not expressing Wor-GAL4. **F)** Control and *flare* RNAi-expressing neuroblasts expressing Cindr::GFP. The arrows point to a neuroepithelial region not expressing *worniu*-GAL4. **G)** Two scenarios could explain the mispositioning of Baz crescents in interface components-depleted neuroblasts: an abnormally mobile centriole directing polarization at another place, or the polarizing effect of the NB/GMC interface is inactive. Measuring the displacement of the apical centriole from polarization (as soon as neuroblasts start rounding up) to metaphase (when the centriole always relocalizes to the center of the apical crescents) allows to discriminate between these possibilities. Centriole position in prophase was not considered, as centrioles transiently detach from the cortex at that phase. **G’)** Displacement of the centriole explained in G), in control (1.7±0.7µm n=75) and *flare* RNAi-expressing neuroblasts (2.2±1.0µm, n=83). The significantly higher displacement in Flare-depleted neuroblasts supports the “disrupted polarizing cue” scenario.

## Movie captions

**VIDEO 1.** Three successive divisions of a neuroblast expressing Worniu-GAL4-driven PH::GFP, shown in **Figure S1A**. Middle panel: orthogonal slice along the y axis of the left panel. Right panel: 3D reconstruction of the neuroblast volume, sliced in half through its division axis. Time in hh:mm:ss, scale bar: 5µm.

**VIDEO 2.** Two successive divisions of a neuroblast expressing PH::RFP (magenta), the centriole marker Asl::YFP and the apical polarity marker Baz::GFP (green), shown in **Figure S1C**. Baz::GFP starts accumulating apically as the neuroblast starts rounding up, around 00:30:00. Arrow: apical centriole. Arrowheads: limits of the apical Baz crescent. Time in hh:mm:ss, scale bar: 5µm.

**VIDEO 3.** Immediate deformation of a neuroblast expressing Worniu-GAL4-driven PH::GFP, following the ablation of its last daughter cell (occurring at 0s, red lightning symbol). Scale bar: 5µm

**VIDEOS 4 & 5.** Two successive divisions of neuroblasts expressing Worniu-GAL4-driven PH::GFP, following control (VIDEO 4) or GMC (VIDEO 5) ablation (red lightning symbol), shown in **Figure 2A**. Arrows: division axis. Time in hh:mm:ss, scale bar: 5µm.

**VIDEO 6.** Two successive divisions of a neuroblast expressing Worniu-GAL4-driven PH::GFP, ending after the formation of plasma membrane extensions at the neuroblast/last-born GMC interface, followed by a high-resolution series of confocal slices through the neuroblast volume, followed by a 3D reconstruction of this volume. Time in hh:mm:ss, scale bar: 5µm.

**VIDEO 7.** Neuroblast expressing Cindr::GFP at endogenous levels. Cindr::GFP localizes to the neuroblast/GMC interface, to the plasma membrane extensions originating from it and to centrosomes.

**VIDEO 8.** Two successive divisions of a neuroblast expressing PH::RFP (magenta), the centriole marker Asl::YFP, the apical polarity marker Baz::GFP (green), and Worniu-GAL4-driven *cindr* RNAi, shown in **Figure 5B**. Following the first division, the apical centrosome maintains its position throughout interphase, a Baz crescent forms at an abnormal location during cellular rounding, the spindle properly aligns with this crescent and the neuroblast divides with a 53° deviation from the first division axis. Arrow: apical centriole. Arrowheads: limits of the apical Baz crescent. Time in hh:mm:ss, scale bar: 5µm.

